# Universal Transformer-Based Tracker for Accurate Tracking of Particles and Cells in Microscopy

**DOI:** 10.1101/2025.05.09.653028

**Authors:** Yudong Zhang, Dan Liu, Ge Yang

## Abstract

Accurate tracking of subcellular structures and cells under microscopy supports heavily the studies of their dynamic processes. However, the complex motion and the similar appearances of objects pose significant challenges in accurately identifying the identical object across multiple detection results without introducing ambiguities in trajectory tracking. Here we propose a Universal Transformer-Based Tracker (UTT) that achieves the accurate tracking in various biologically relevant scenarios. The self-attention of Transformer extracts features and patterns from trajectories, and the cross-attention of Transformer captures matching relations between history tracks and future hypothesis tracklets (fragments of track). Our tracker uses the Transformer to handle the tracking of particles with diverse motion dynamics, and outperforms existing tracking algorithms in terms of accuracy. Furthermore,the tracker utilizes estimated particle positions to counteract missed detections, thereby improving tracking robustness under varying ratios of missing detections. We demonstrate the flexibility and versatility of our approach by adaptively integrating motion and appearance cues in cell tracking applications.

## 1 Main

The movement trajectories of cells and organelles contain dynamic information about their life activities and reveal many laws in life sciences. For example, May et al.[1] tracked the spinal cord precursor cells of mice and their division behaviors, and reported for the first time the impact of the extracellular matrix glycoprotein Tnc on the cell cycle length of spinal cord precursor cells. Lovas et al.[2] tracked the activities of hydra neurons and revealed the self-organization ability and modular characteristics of hydra in the reorganization of the nervous system, providing important insights for understanding the recovery mechanisms of organisms after suffering injuries. To quantitatively analyze the activity states of these objects, such as characterizing the velocity of object motion and the cell division cycle, accurate tracking of the objects’ movement trajectories is essential. Therefore, tools for single-particle tracking and single-cell tracking are of paramount importance.

Most single-particle tracking and single-cell tracking methods follow the tracking-by-detection(TBD) paradigm[3–5], which consists of two steps: (1) For each frame of the time-lapse imagery captured by microscopes, detectors are used to obtain the localization information or mask information of all objects; (2) By measuring the degree of match between detection results, all detections are reasonably linked to form several trajectories.

Based on the size of the objects, microscopic objects can be categorized into sub-cellular structures and cells. Subcellular structures are typically smaller than the diffraction limit of optical microscopes, thus exhibiting the characteristics of an Airy disk. Their numerous similar appearances and complex and variable motion patterns pose challenges to the tracking task. Additionally, their small size makes them easily missed in detection, leading to inaccuracies and discontinuities in the reconstructed trajectories. Cells are larger and possess a richer set of morphological features than subcellular structures. Additionally, cells exhibit unique division behaviors. Therefore, it is necessary not only to identify the trajectories of the same target but also to correctly recognize the mother-daughter relationship of dividing cells. Due to the different characteristics of the subcellular structures and cells, tracking methods for these two types of targets are designed and developed separately.

When tracking particles, most conventional methods rely primarily on motion information for association. These methods[6–9] are based on the assumption of a deterministic motion pattern, thus a single approach may not be flexible enough to handle complex motion behaviors. Moreover, these methods often require manual tuning of model parameters and cannot track automatically. Recently, deep learning-based tracking algorithms[10–12] have emerged, utilizing recurrent neural networks and fully connected networks to learn trajectory motion features and predict associations, achieving automatic tracking. Recent advanced cell tracking methods mostly use appearance and morphological features for association[13–15]. However, they do not take into account the varying degrees of appearance specificity among different types of cells. In cases where the target cells have similar appearances, the performance of these methods may deteriorate or even fail.

In order to automate the tracking of diverse targets including various particles and cells, we conceived the idea of adaptively leveraging motion information and appearance information. By flexibly adjusting the degree of dependence on these two types of features, we can fulfill the tracking requirements of different target types. Specifically, we proposed a universal tracking method based on the Transformer, namely UTT. The self-attention mechanism is utilized to extract the temporal information of trajectories, and the cross-attention mechanism is employed to capture the matching relationships between historical trajectories and future hypothesized trajectory segments. Concerning the missed detections of particle targets, a reconnection strategy is devised within our method. By using network predictions to identify missed detection scenarios and leveraging the estimated positions to compensate for the missed detections, our method can effectively handle such situations. We adopted two branches to separately utilize the motion information and the appearance information and then applied a gating mechanism to adaptively fuse these two types of cues to adapt to the tracking demands of different target types. We have conducted extensive experiments on particle data with various motion characteristics and data of different cell types, and the experimental results have validated the superiority of our method in both particle tracking and cell tracking.

## 2 Results

### 2.1 Dual-experts capture matching relations

The Universal Transformer Tracker (UTT) adopts a detection-based tracking paradigm, that operates in a detect-then-track manner. The detection step utilizes the deepBlink algorithm[16] to identify particle positions in each frame. For tracking step, the system initializes trajectories in the first frame, then data association and trajectory state updates are iteratively executed for each frame until the end of the video, as shown in Fig.1.

It is assumed that the trajectories up to frame *t* have been completed, and the association between frame *t* and frame *t* + 1 needs to be carried out. Fig. 1a shows detection results on three consecutive frames (t, t+1, t+2). Then, we select candidate trajectories for each historical trajectory across multiple future frames. For example, as shown in Fig. 1b, for the historical trajectory *H*_*i*_, candidate points are identified from the detection points in future frames based on the Euclidean distance metric. Details see Methods 4.2. For each historical trajectory, a set of candidate short trajectories is generated, as illustrated in Fig. 1c (e.g., *C*_*k*_, *k* = 1, 2, …, 9). Subsequently, motion features from historical trajectories and candidate trajectories, as well as appearance features (only for cells), are extracted. These trajectory features are then input into a transformer network to further refine the matching relationships, as shown in Fig. 1d. The model predicts the matching probabilities between each historical trajectory and its candidates, and then are aggregated to the *t* + 1 frame’s detections via max-polling. Meanwhile, the model predicts the future coordinates of historical trajectories and their existence scores (i.e., the probability of the trajectory not terminating). Subsequently, leveraging the matching probability matrix between frames *t* and *t* + 1 (Fig. 1e), we formulate a global optimization equation to obtain the optimal solution that satisfies the constraints (Fig. 1f). Following this, we manage the trajectories based on both the outcomes of the global optimization and the predicted future states of the trajectories from the network. Particularly, for those results matched to null detections, we contemplate performing reconnection operations. The details is described in the Methods section and Supplementary Materials, Fig. B2.

**Fig. 1.**
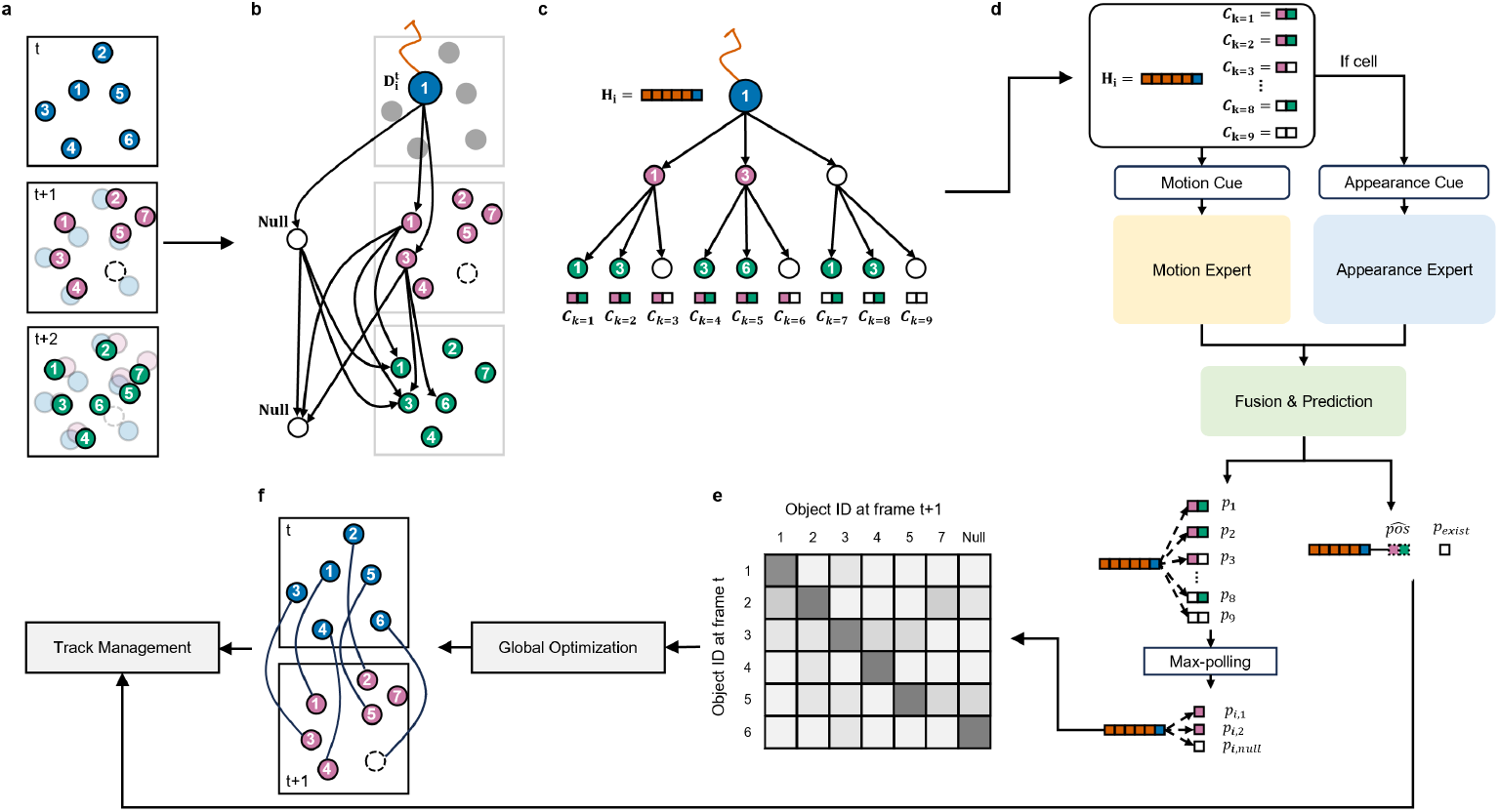
Pipeline of our method UTT. a. Detection results of targets on three consecutive frames (t, t+1, t+2) are illustrated, with different colors used to distinguish results from different frames. The numbers within the circles denote the IDs of the corresponding targets. On frame t, six detections are present. On frame t+1, target 6 is undetected, while a new target 7 emerges. On frame t+2, target 6 reappears. b. A trajectory that concludes with detection 1 on frame t is depicted. Candidate detections are chosen based on distance for the subsequent two frames. “Null” signifies the absence of a detection in future frames. c. The selected detections construct a hypothesis tree, yielding a total of nine candidate short trajectories. d. Matching probabilities and future state prediction network. The matching probabilities between historical trajectories and detections in the next frame, as well as the future states of historical trajectories, are derived. e. The predicted matching probability matrix is displayed, with darker shades of gray indicating higher matching probabilities. f. The optimal solution that satisfies the constraints is determined through global optimization.

We evaluate our method on the Receptor scenario of ISBI particle tracking dataset (as shown in Fig.C3), achieving 97.7% frame-to-frame matching accuracy and 88.9% prediction accuracy within 5 pixels. Figure D4a-d show particle tracking in dense regions (snr 7, high density), where our model correctly matches particles despite near-identical appearances (Fig. D4f) and handles disappearing particles (e.g., particle 6). Trajectory predictions achieve 2.04 pixels average error, with Fig. D4g-h demonstrating consistent accuracy even in crowded regions. These results demonstrate the effectiveness of our approach in achieving accurate matching and trajectory prediction, even in complex scenarios involving dense particle arrangements, occlusions, and high appearance similarity.

### 2.2 UTT tracking particles accurately

The performance of our method was comprehensively evaluated on the ISBI Particle Tracking Challenge (PTC) dataset (refer to Fig.C3) across diverse scenarios and conditions, as summarized in Fig. 2. Fig. 2**a** demonstrates the method’s accuracy under ground truth detection conditions across four biological scenarios (Microtubules, Vesicles, Receptors, Viruses) and three particle densities (Low, Middle, High). Consistently high performance is observed, with the microtubules scenario showing the best results, highlighting the method’s adaptability to varied particle dynamics and densities.

**Fig. 2.**
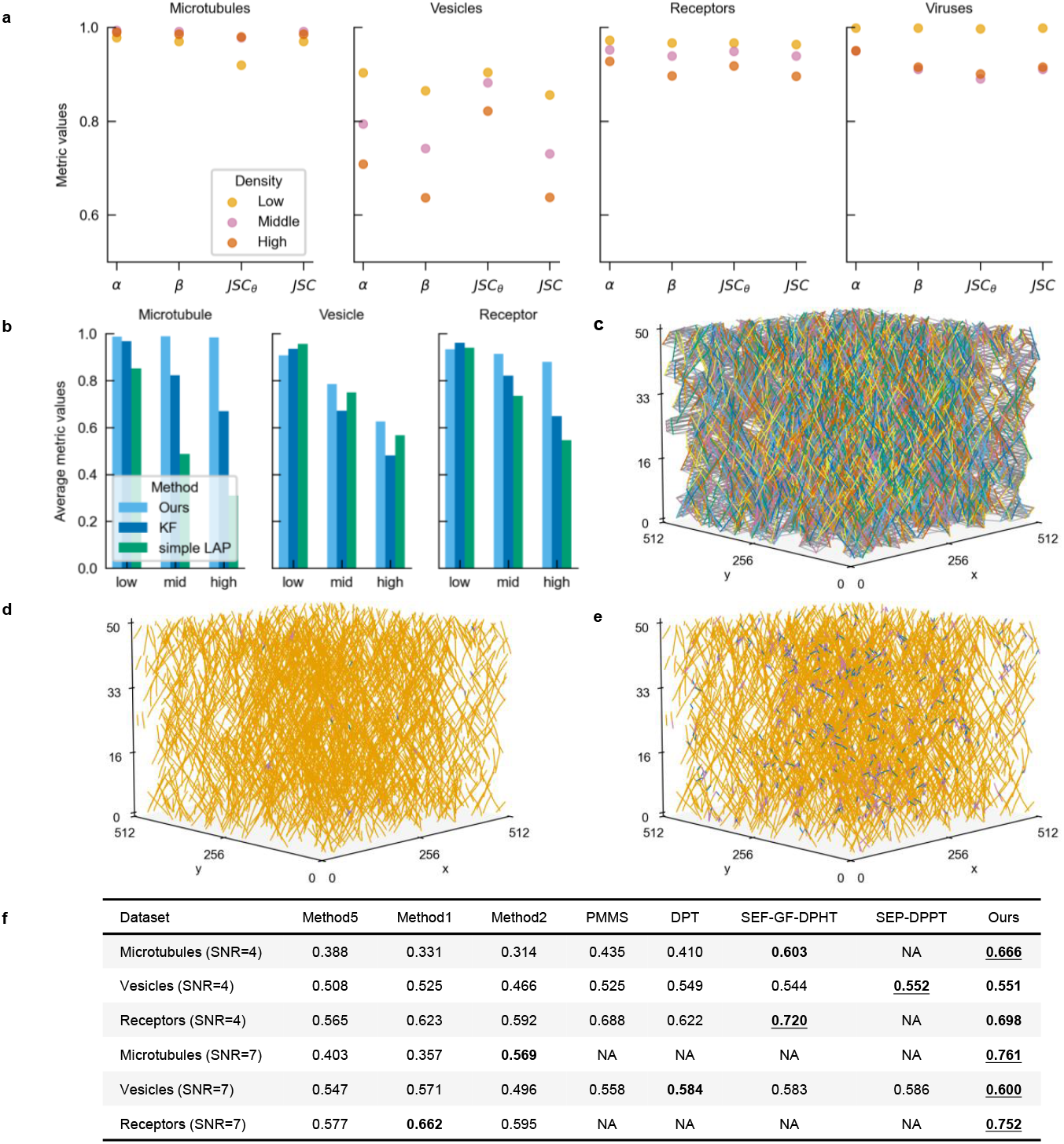
Particle tracking results on ISBI PTC dataset. **a**, Performance evaluation on ISBI PTC test data across four scenarios (Microtubules, Vesicles, Receptors, Viruses) and three densities, showing robust performance with best results in Microtubules. **b**, Comparative analysis with traditional methods (KF, sampleLAP) using ground truth detections. While all methods perform well (metrics *>*0.9) at low density, our method outperforms at middle/high densities across the average of four metrics (*α, β, JSC*_*θ*_, *JSC*, details see Methods 4.7) in 2D scenarios. **c-e**, Tracking visualization on Microtubule data (SNR7, mid-density, frames 0-50): **c**, shows ground truth trajectories (colored) versus hypothetical connections (gray); **d**,**e**, compare our method (fewer errors) with KF, using color-coded results (correct: orange, over-linking: blue, under-linking: pink). **f**, Benchmark against SOTA methods using their detections on high-density data (four metrics averaged). Our method achieves top performance (bold/underlined = 1st place, bold = 2nd place; NA = unreported metrics).

Fig. 2**b** compares our method with traditional algorithms such as the Kalman Filter (KF) and sampleLAP under identical ground truth detection settings. While all methods perform well at low particle density, our method demonstrates significantly better performance at medium and high densities, showcasing its robustness in handling complex scenarios like dense clustering.

To further assess the accuracy of trajectory recovery, Fig. 2**c-e** provide detailed comparisons of trajectory connections for the MICROTUBULE video under mid-density with directional motion. Fig. 2**(c)** shows ground truth trajectories (colored) versus hypothetical connections (gray). It can be observed that the trajectories are densely distributed with numerous intersections. Figs. 2**(d)** and **(e)** present the tracking results of our method and the Kalman Filter (KF), respectively. Correct connections matching the ground truth are marked in orange, over-connections in blue, and missed connections in pink. The results demonstrate that our method achieves accurate tracking with minimal errors, as evidenced by fewer false connections (over-linking, blue lines) and missed connections (under-linking, pink lines), compared to the Kalman Filter results in Fig. 2**e**.

Finally, Fig. 2**f** evaluates our method against other state-of-the-art (SOTA) methods using their respective detection results on high-density data. Across key metrics (*α, β, JSC*_*θ*_, *JSC*), our method ranked first on four datasets (Microtubules at SNR=4/7, Vesicles and Receptors at SNR=7) and second on two others (Vesicles and Receptors at SNR=4). Consequently, our method exhibits optimal performance in comprehensive evaluation. Moreover, this comprehensive comparison emphasizes the superior accuracy and robustness of our approach in addressing the challenges posed by high-density particle tracking scenarios.

Together, these results demonstrate that our method effectively integrates robust tracking algorithms and accurate data association strategies, achieving state-of-the-art performance across diverse and challenging biological tracking tasks.

### 2.3 Re-linking strategy improves tracker robustness

To enhances the robustness of particle tracking, in scenarios where false negatives (FN) can disrupt trajectory continuity, we propose the re-linking strategy. As shown in Fig. 3a, for historical trajectories matched to empty in the global optimization results, we do not directly terminate them. Instead, we further evaluate the predicted existence probability score of these trajectories. If the score exceeds a predefined threshold, we determine that the trajectory should not be terminated but rather represents a missed detection. Consequently, we utilize the model’s predicted future coordinates of these historical trajectories to fill the missed detections, thereby preventing trajectory fragmentation. The strategy systematically addresses tracking errors by leveraging global optimization results and existence probabilities to correct mismatches and maintain accurate associations across consecutive frames.

**Fig. 3.**
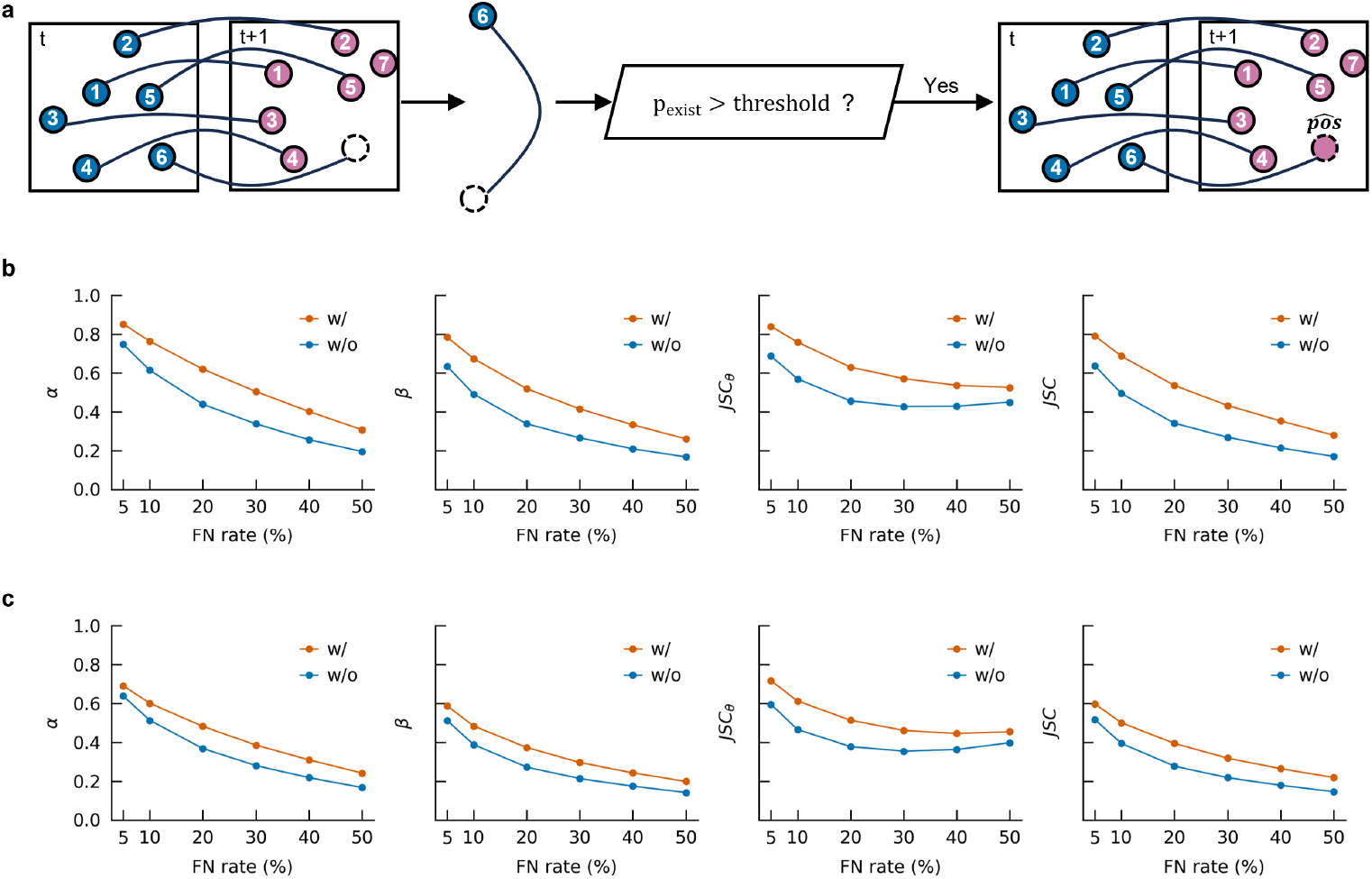
Re-linking strategy flowchart and quantification comparison. **a**, Re-linking strategy. For cases in the global optimization results where matches are made to null detections, we assess whether the likelihood of their existence exceeds a predefined threshold. If the likelihood surpasses this threshold, the case is considered a missed detection. We then utilize the model-predicted coordinates to fill in for the missing detection. **b**,**c**, Comparison of quantification results with(w/) and without(w/o) the re-linking strategy across two particle scenarios at various false-negative (FN) ratios (5%, 10%, 20%, 30%, 40%, 50%). **b**, Comparison at medium density in the receptor particle scenario. **c**, Comparison at medium density in the vesicle particle scenario.

To evaluate the effectiveness of this strategy, experiments were conducted across different particle scenarios at various false-negative (FN) ratios. The quantification results, shown in Fig. 3b,c, demonstrate that the re-linking strategy significantly improves tracking performance in receptor and vesicle particle scenarios under varying FN conditions.

Visual evidence provides further validation of the re-linking strategy’s effectiveness. In the receptor scenario (Fig. F6a-c), false negatives at different frames lead to changes in track IDs without applying the re-linking strategy. However, the re-linking strategy ensures trajectory continuity by reconnecting trajectories based on their existence probabilities, resulting in a closer alignment with the ground truth trajectories. This visual comparison (Fig. F6a-c) underscores the strategy’s ability to effectively mitigate tracking errors and maintain the accuracy of particle associations across frames.

Overall, the re-linking strategy significantly enhances the robustness of particle tracking by addressing false negatives, improving tracking accuracy, and ensuring continuous trajectories.

### 2.4 Integrating motion and appearance cues improving cell tracking performance

Compared to particle-like objects, cell-like targets exhibit larger sizes and distinct visual characteristics, such as texture and morphology. To enhance tracking performance for such targets, we propose to integrate motion and appearance features for association. Specifically, we evaluate four feature utilization strategies: M (motion features only), M+MA (motion features with handcrafted appearance features), M+CA (motion features with CNN-based appearance features), and M+MA+CA (motion features with both handcrafted and CNN-based appearance features). As shown in Fig. 4. M+CA significantly outperformed M (*p <* 0.05) in terms of the AOGM metric (Fig. 4**a**), primarily due to a marked reduction in false negative edges (*fn_edges*), as shown in Fig. 4**b**. This indicates that appearance features effectively address missed connections, improving tracking accuracy.

**Fig. 4.**
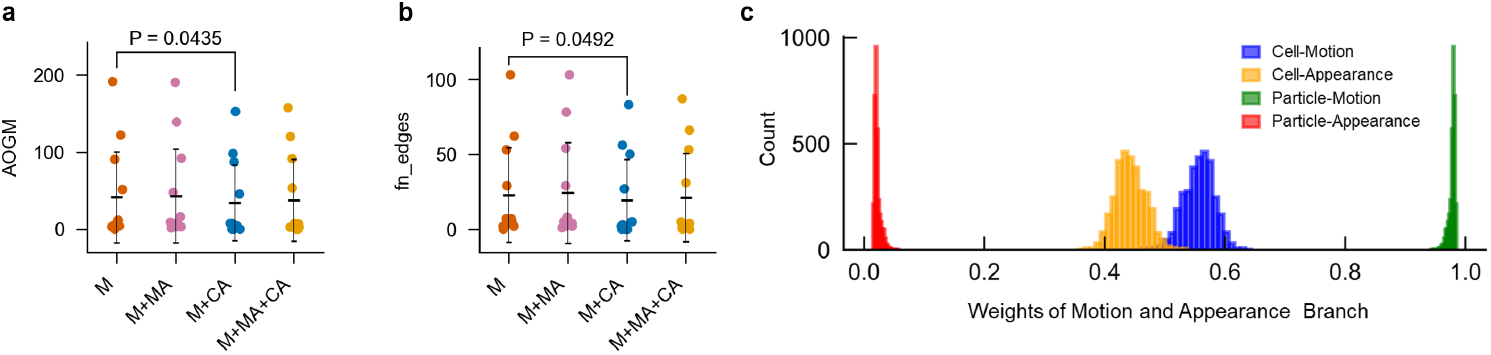
Impact of integrating motion and appearance features on cell tracking performance. **a-b**, Performance metrics under four feature configurations: M (motion features only), M+MA (motion features combined with handcrafted appearance features), M+CA (motion features combined with CNN-based appearance features), and M+MA+CA (motion features combined with both handcrafted and CNN-based appearance features). **a**, Results for the AOGM metric. **b**, Results for the fn edges metric. **c**, reliance of UTT on appearance and motion information when tracking particles that are visually similar as well as cells with distinct appearances.

Given the differing levels of appearance specificity between cells and particles, we propose an adaptive fusion mechanism to improve the model’s universality across both target types. Our approach employs two parallel branches: one for motion-based matching and another for appearance-based matching, coupled with a gating network that dynamically adjusts the reliance on each branch based on target characteristics, details see B2. As illustrated in Fig. 4c, the model autonomously adapts its tracking strategy: when tracking particles that are visually similar, the model assigns a weight close to 1 to the motion branch, primarily relying on motion information for tracking. In contrast, when tracking cells with distinct appearances, the model utilizes both motion and appearance information for tracking. This adaptive model design significantly enhances the versatility of our method.

## 3 Discussion

Accurate tracking of particles and cells is crucial for quantitative analysis in biological microscopy data. However, this task faces significant challenges due to the similar appearance, high density, and complex motion patterns of particle targets. Additionally, the varying specificity levels of cellular and particulate features make it difficult to establish a unified tracking approach. Here, we introduced a Universal Transformer-Based Tracker, UTT, addresses these challenges by adaptively leveraging both motion cues and appearance cues, enabling universal tracking of targets with diverse feature specificity. UTT employs self-attention to effectively extract temporal features and cross-attention to model matching relationships. Compared with fully connected networks, the cross-attention mechanism can more effectively predict the degree of matching. Furthermore, we introduce a re-linking strategy to recover missed detections by predicting future coordinates, which enhances robustness in tracking continuity. Experimental results demonstrate that our model dynamically adjusts its reliance on appearance and motion cues depending on the target’s distinctiveness. Moreover, UTT achieves state-of-the-art performance across multiple scenario, particularly excelling in high-density, challenging scenarios. Future work may explore large-scale training within this framework to further improve generalization.

## 4 Methods

### 4.1 Pipeline of tracking

The proposed particle tracking method follows a two-step framework: particle detection and particle linking. As shown in Fig. A1, first, we use the deepBlink detector to detect particles in each frame. The detections in the first frame are initialized as active tracks. In subsequent frames, we perform particle linking through four main steps. Firstly, for each active track, we construct a hypothesis tree to generate multiple future hypothetical tracks. Then, all tracks, after preprocessing, are input into the UTT network, which predicts the matching probabilities between each active track and its multiple future hypothetical tracks, as well as the probability and position of each active track in the next frame. Subsequently, a discrete optimization model is established to find the globally optimal matching for all active tracks by maximizing the total matching probability. Finally, a track management strategy is designed to handle track initialization, updating, termination, and re-linking. Overall, by iteratively matching historical tracks to candidate detection points and updating tracks frame by frame, the method accurately recovering the trajectories of all particles in the video.

### 4.2 Construction of hypothesis trajectories

To establish the correspondence between history trajectories and detections in frame (*t* + 1), multiple hypothesis candidate trajectories are generated for each historical trajectory by constructing a hypothesis tree. For a given historical trajector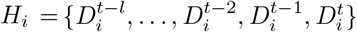, a hypothesis tree of depth *d* is built, where each non-leaf node has *m* child nodes. The last detection point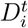 serves as the root of the tree. The tree is expanded by selecting candidate detections based on Euclidean distance.

Specifically, for each node, the (*m−* 1) closest detections in the next frame are selected as children, along with one Null detection representing a potential missed detection. If the current node is a Null node, the (*m −*1) closest detections are instead selected based on the distance to its parent node. In practice, the tree is constructed using a breadth-first search algorithm. This results in a total of *K* = *m*^*d*^ candidate trajectories. In this way, our method considering multiple frames hypothesis when predicting matching relations between history trajectories and the next frame detections. Fig. 1 shows an example of constructing hypothesis trajectories for a particle with *m* = 3 and *d* = 2.

### 4.3 Prediction of Matching Probability and Future States

To enhance the accuracy and universality of association prediction in multi-object tracking tasks, we propose a trajectory matching model based on adaptive feature fusion, as illustrated in Fig. **??**. The model consists of a motion branch and an appearance branch, which respectively model the association likelihood based on motion features and appearance features. A gating mechanism is then used to dynamically allocate weights between the two branches, and the final matching probability is produced by fusing their predictions.

The motion branch comprises a motion encoder, a Transformer-based temporal feature and matching extractor, and two output heads. The inputs include the coordinate sequences of historical trajectories and all candidate trajectories. An additional binary dimension *f* is introduced to indicate the validity of each detection, where *f* = 1 denotes a real detection and *f* = 0 denotes a dummy detection. Thus, for 2D trajectories, the input format is (*x, y, f*).

The motion encoder first computes displacement features by differencing the coordinate sequences and adjusts the validity dimension accordingly. For the first coordinate of each candidate trajectory, displacement is calculated relative to the last coordinate of the historical trajectory. These features are then embedded via a fully connected network. The resulting motion embeddings have shape (*B, L, c*) for historical trajectories and (*B, K, d, c*) for candidate trajectories, where *B* is the batch size, *L* is the length of the historical trajectory, *K* is the number of candidates, *d* is the candidate trajectory length, and *c* is the embedding dimension.

The motion embeddings are fed into a Transformer to extract temporal features and matching relationships. The historical trajectory embedding, combined with positional encoding, is passed through a Transformer encoder that captures global temporal dependencies via self-attention and further refines the features using a feedforward network (FFN). For candidate trajectories, the temporal and embedding dimensions are merged to form an input of shape (*B, K, d ×c*), which is then processed by a Transformer decoder. The decoder first applies self-attention to enhance internal representations and then performs cross-attention using encoder outputs as keys and values, and decoder inputs as queries. This captures the association between each candidate and the historical trajectory. The output of the decoder is a matching feature vector of shape (*B, K, c*_1_), which is used by a classification head to predict motion-based matching scores *P*_*m*_ *∈* ℝ^*B×K*^. A regression head is also used to predict future positions of the target, which facilitates trajectory reconnection in particle tracking.

The appearance branch consists of an appearance feature encoder, a Transformer-based feature and matching extractor, and a classification head. The inputs are sequences of cropped image patches centered on each target in the historical and candidate trajectories, with size *W× H*. The image patches are passed through a convolutional neural network (CNN) to extract low-level features, which are then flattened and embedded using a fully connected network to produce appearance embeddings. These embeddings are passed through the Transformer encoder-decoder architecture similar to the motion branch. The decoder output has shape (*B, K, c*_1_) and is used to compute the appearance-based matching scores *P*_*a*_ *∈* ℝ^*B×K*^ via a classification head.

A gating network adaptively fuses information from the two branches by assigning dynamic weights. Specifically, the motion and appearance matching vectors are aggregated across candidate trajectories to obtain two global feature vectors. These are concatenated and passed through a multi-layer perceptron (MLP) to compute two weights *w*_*m*_ and *w*_*a*_, which adjust the contributions of the motion and appearance branches. The final matching probability is obtained by weighted fusion and softmax normalization:

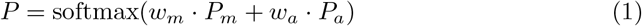

where *P* ∈ ℝ^*B×K*^.

During training, we ensure that the candidate set of each historical trajectory contains one ground truth trajectory. The predicted matching probabilities and predicted future states are supervised using cross-entropy loss and mean squared error (MSE) loss respectively.

During inference, we apply max pooling across candidate trajectories to obtain the final matching probability *P*_0_ for each detection in the next frame, with shape (*B, m* + 1), where *m* is the number of detections in the current frame. The pooling operation is defined as:

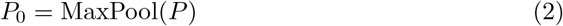

### 4.4 Discrete optimization problem modeling

After obtaining the predicted matching probability distribution between historical tracks and next-frame detections, we formulate the assignment as a discrete optimization problem. Specifically, we aim to find a feasible one-to-one correspondence that maximizes the overall matching confidence while satisfying matching constraints.

Let *p*_*ij*_ *∈* [0, 1] denote the predicted matching probability between active track *i* and detection *j*, and let *a*_*ij*_ ∈{0, 1} be a binary decision variable indicating whether track *i* is assigned to detection *j*. In particular, *j* = 0 represents a dummy (null) detection, allowing for unmatched tracks.

The objective is to maximize the total matching score subject to assignment constraints: each active track must be assigned to exactly one detection (including the dummy), and each real detection can be assigned to at most one track. The optimization problem is formulated as:

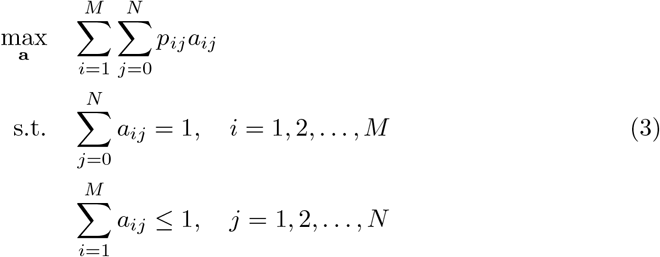

This optimization problem is solved using the CBC (Coin-OR Branch and Cut) solver [17], which is an open-source mixed-integer programming solver. CBC is designed to solve large-scale linear and integer programming problems by using branch-and-cut techniques, which combine branching, cutting planes, and other advanced algorithms to efficiently explore feasible solutions. It is widely used in various optimization applications due to its flexibility and performance.

### 4.5 Tracks management

To maintain consistent target identities across frames and handle issues such as missed detections and false positives, trajectory management is performed after obtaining the one-to-one matching results. This strategy enables effective identification of trajectory initialization, updating, and termination. Specifically, based on the optimization results, history trajectory segments that are matched to true detections are updated by appending the matched detection. Segments matched to null detections are either terminated or updated with predicted positions, depending on whether the predicted existence probability exceeds a predefined threshold *p*. This mechanism—referred to as the relinking strategy—allows broken trajectories caused by missed detections to be preserved and potentially reconnected when the targets reappear. Meanwhile, unmatched detections are used to initialize new history trajectory segments. After tracking particles throughout the sequence, trajectories of length one are removed, as they are considered false positives. Notably, particle targets require this relinking strategy to address the common occurrence of missed detections, while cell targets—due to their larger size—are less prone to detection failures and therefore do not require relinking. For a comprehensive explanation of the procedure, refer to the pseudocode in Algorithm 1.

### 4.6 Lineage reconstruction

Cell division is specifically manifested as a single parent cell in the previous frame splitting into two daughter cells in the next frame. Typically, the parent cell is spatially close to the two daughter cells, but there are significant differences in their appearance features. Therefore, after the tracking algorithm is executed, the parent-child relationship is not identified and matched.

Based on the characteristics of cell division in space and time, this paper constructs a lineage reconstruction algorithm. Daughter cells from the same division event are typically initialized in the same frame and appear spatially close. We leverage this observation to identify sibling pairs by pairing newly initialized trajectories in the same frame using a distance-based metric. The parent cell typically appears spatially close to its two daughter cells, and its trajectory usually ends in the frame immediately before the daughters are initialized. To identify parent-child relationships, we match stopped trajectories in the previous frame with paired daughter cells using a distance metric. In cases where the parent cell’s trajectory continues and merges with one of the daughter cells, leaving the other initialized alone, we further attempt to match such unpaired trajectories with continuous ones to recover potential parent-child links.

### 4.7 Evaluation metric

#### Particle Tracking Metric

Metrics *α, β, JSC*_*θ*_, *JSC* are used to evaluate the method performance[6]. Metric *α* ∈ [0, 1] quantifes the matching degree of ground truth and estimated tracks, while *β in*[0, *α*] is penalized by false positive tracks additionally compared to *α. JSC*_*θ*_ ∈ [0, 1] and *JSC*∈ [0, 1] are the Jaccard similarity coefficient for entire tracks and track points, respectively. Higher values of the four metrics indicate better performance.

The specific computational procedure is as follows: First, the predicted trajectory set is matched with the ground truth to determine correct/incorrect predictions and generate a confusion matrix. The distance between the predicted trajectory set *X* and the ground truth set *Y* is defined as the sum of distances for all matched trajectory pairs:

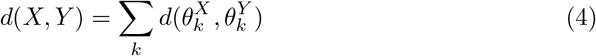

where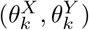 denotes a matched trajectory pair. The distance between each pair is computed as the sum of point-wise distances across all time steps:

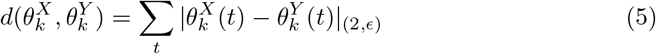

Here, distances exceeding *ϵ* are clipped to *ϵ*. Based on this, the four evaluation metrics are derived as:

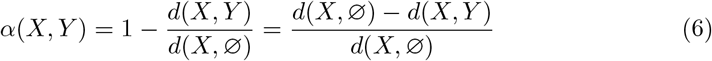

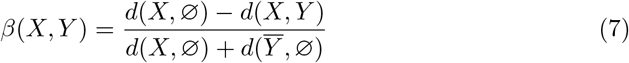

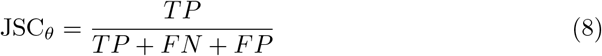

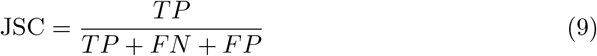

where 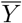 represents false positive (FP) trajectories.

#### Cell tracking Metric

This paper adopts the Directed Acyclic Graph (DAG) metric to evaluate cell tracking performance. The Acyclic Oriented Graph Matching (AOGM) metric operates by converting both tracking results and ground truth (GT) into directed acyclic graphs, then measuring the operational cost required to transform the predicted graph into the GT graph. The specific procedure is as follows:

1. Graph Matching: Align the predicted graph with the GT graph based on detection (segmentation) results.
2. Cost Calculation: Record operational costs during the graph transformation process, which includes:

(1) NS nodes: Count of splitting incorrectly matched predicted nodes into multiple GT nodes. (2) FN nodes: Count of adding missing nodes. (3) FP nodes: Count of deleting redundant nodes and their associated edges. (4) FP edges: Count of deleting redundant edges. (5) FN edges: Count of adding missing edges. (6) WS edges: Count of correcting edge semantics (e.g., linear links vs. division links).

The AOGM score is then defined as the weighted sum of these operational costs, with weights following the Cell Tracking Challenge [**?** ] standards:

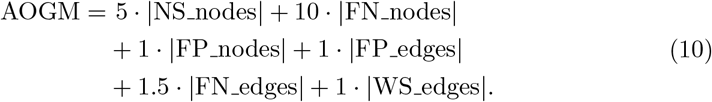

The AOGM metric holistically evaluates tracking quality by accounting for diverse error types, including implicit division events.

Since this study focuses on tracking algorithm design, all experiments use GT segmentation for fair comparison, reducing node-related costs to zero. Thus, we primarily report AOGM, FP edges, FN edges, and WS edges in this paper.

## Appendix A Workflow of the proposed method for multi-object tracking

**Fig. A1.**
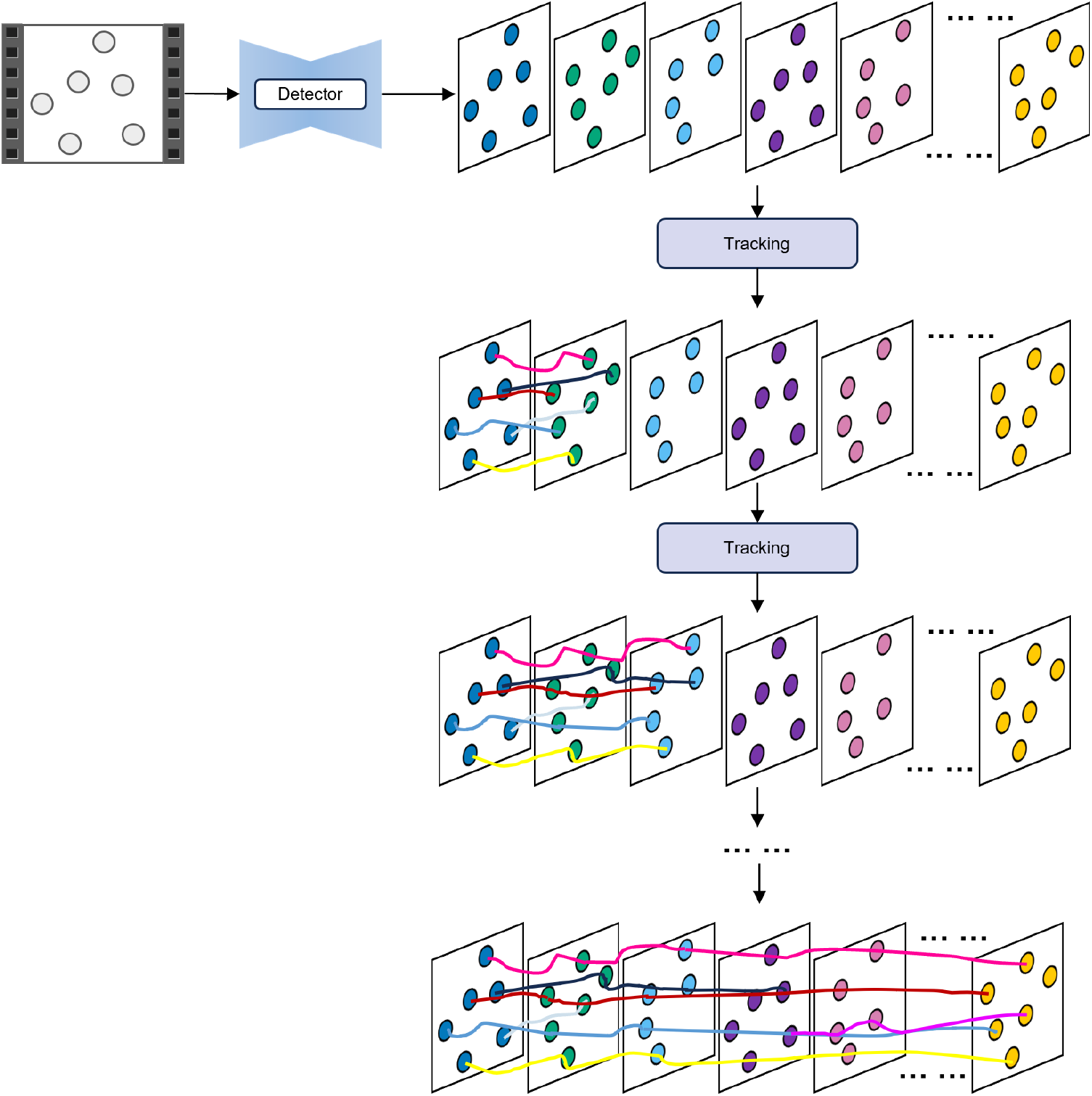
Workflow of the proposed method for multi-object tracking.

## Appendix B Structure of transformer-based network

**Fig. B2.**
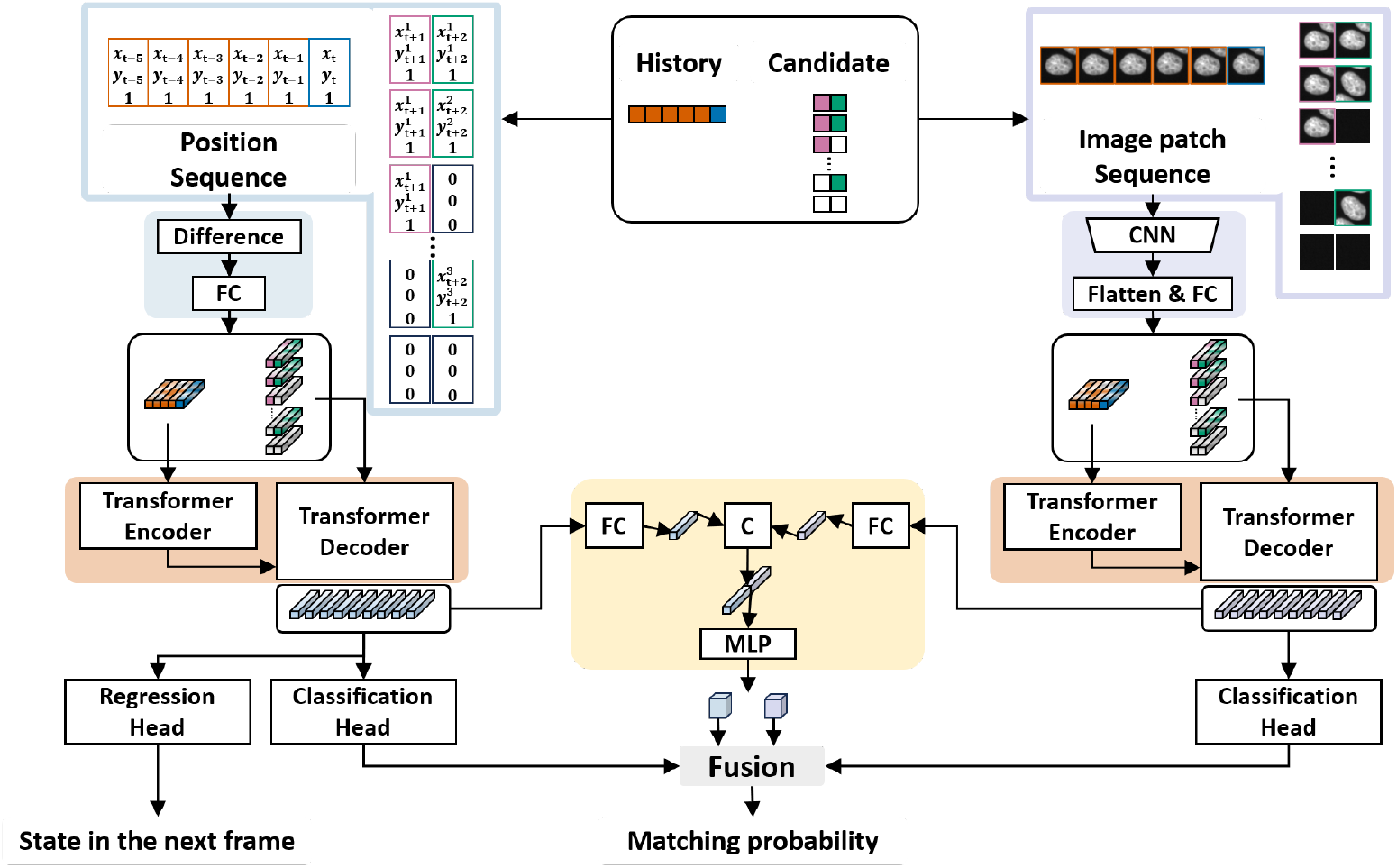
Structure of transformer-based network.

## Appendix C ISBI Particle Tracking Challenge Dataset

**Fig. C3.**
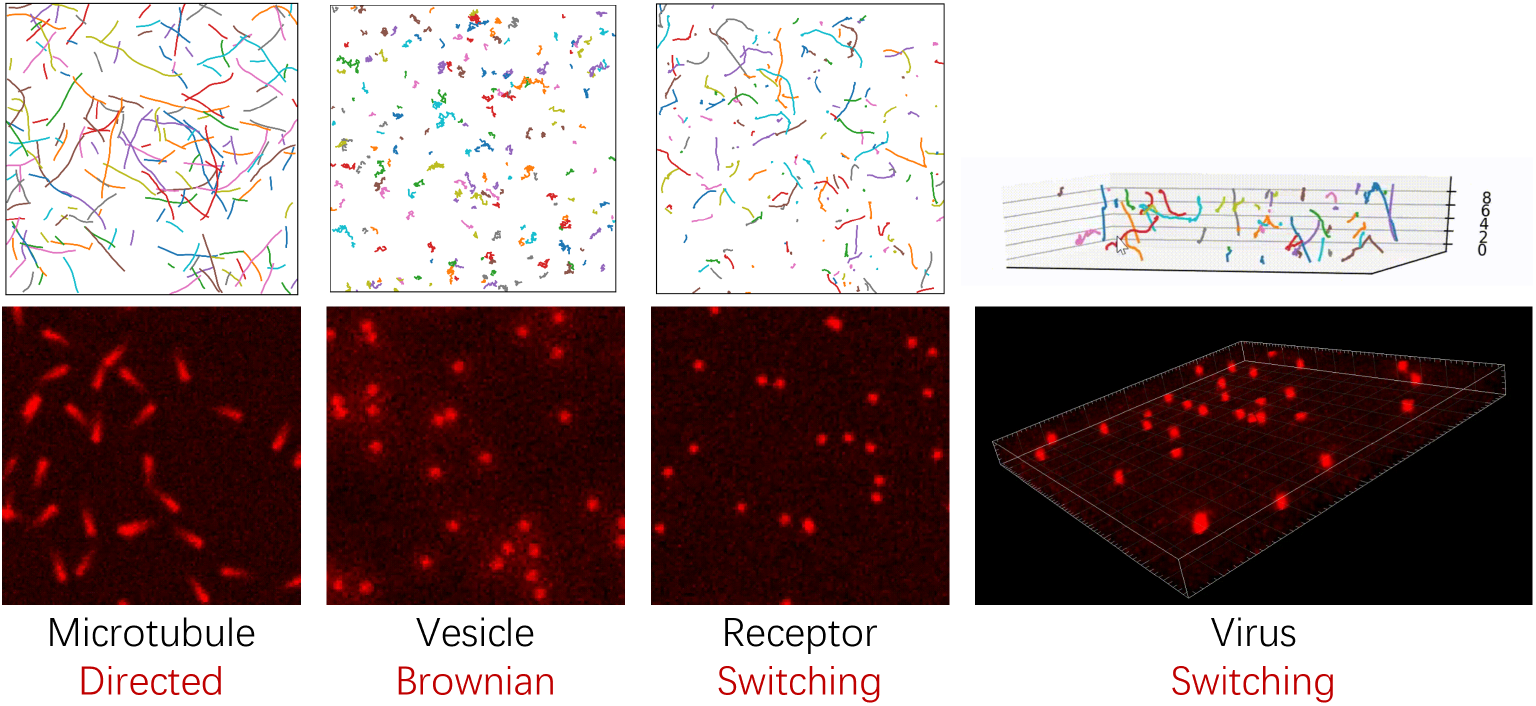
ISBI Particle Tracking Challenge Dataset.

## Appendix D Matching and future position prediction performance

**Fig. D4.**
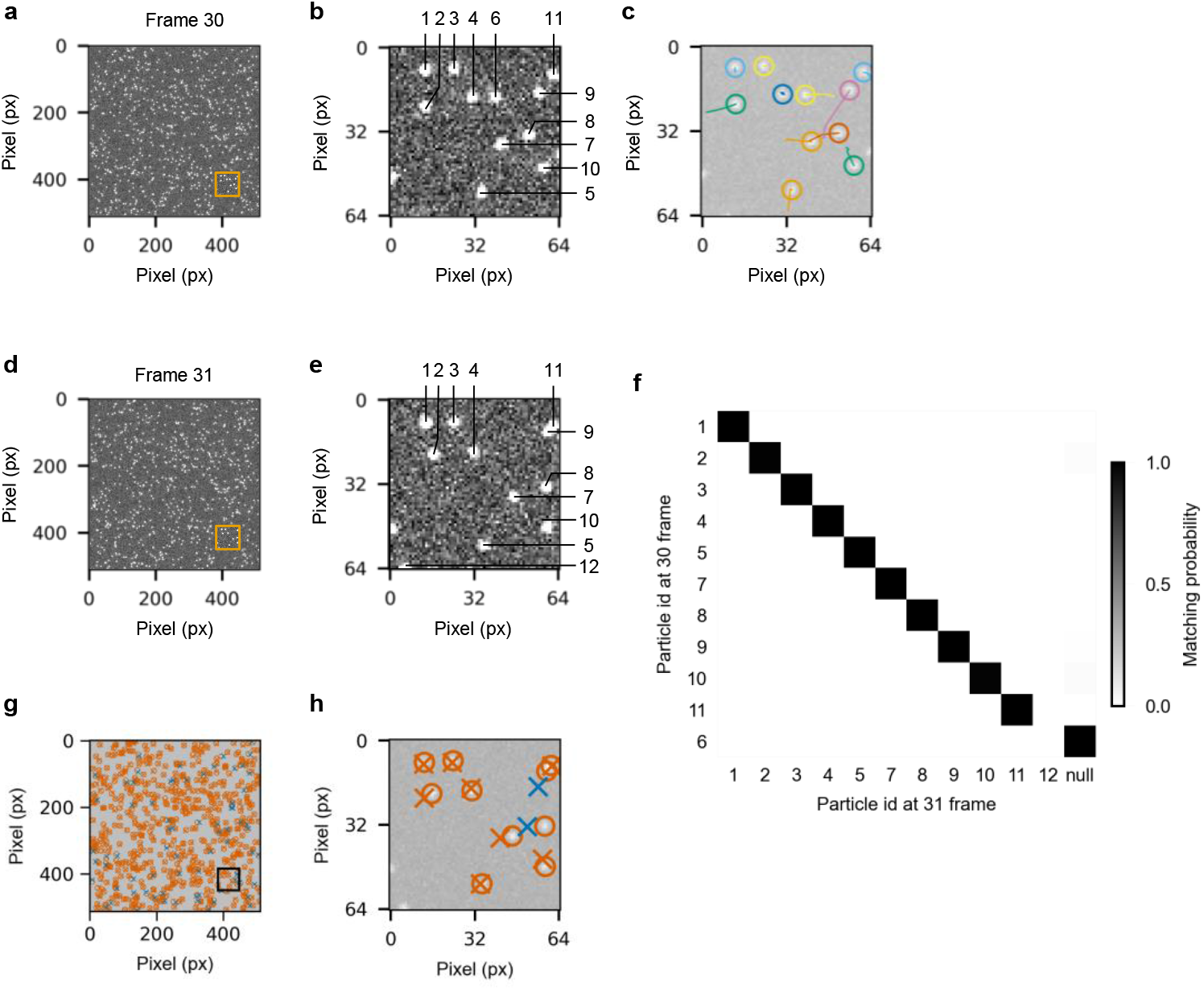
Matching and future position prediction performance.

## Appendix E Particle Tracking performance comparison

**Fig. E5.**
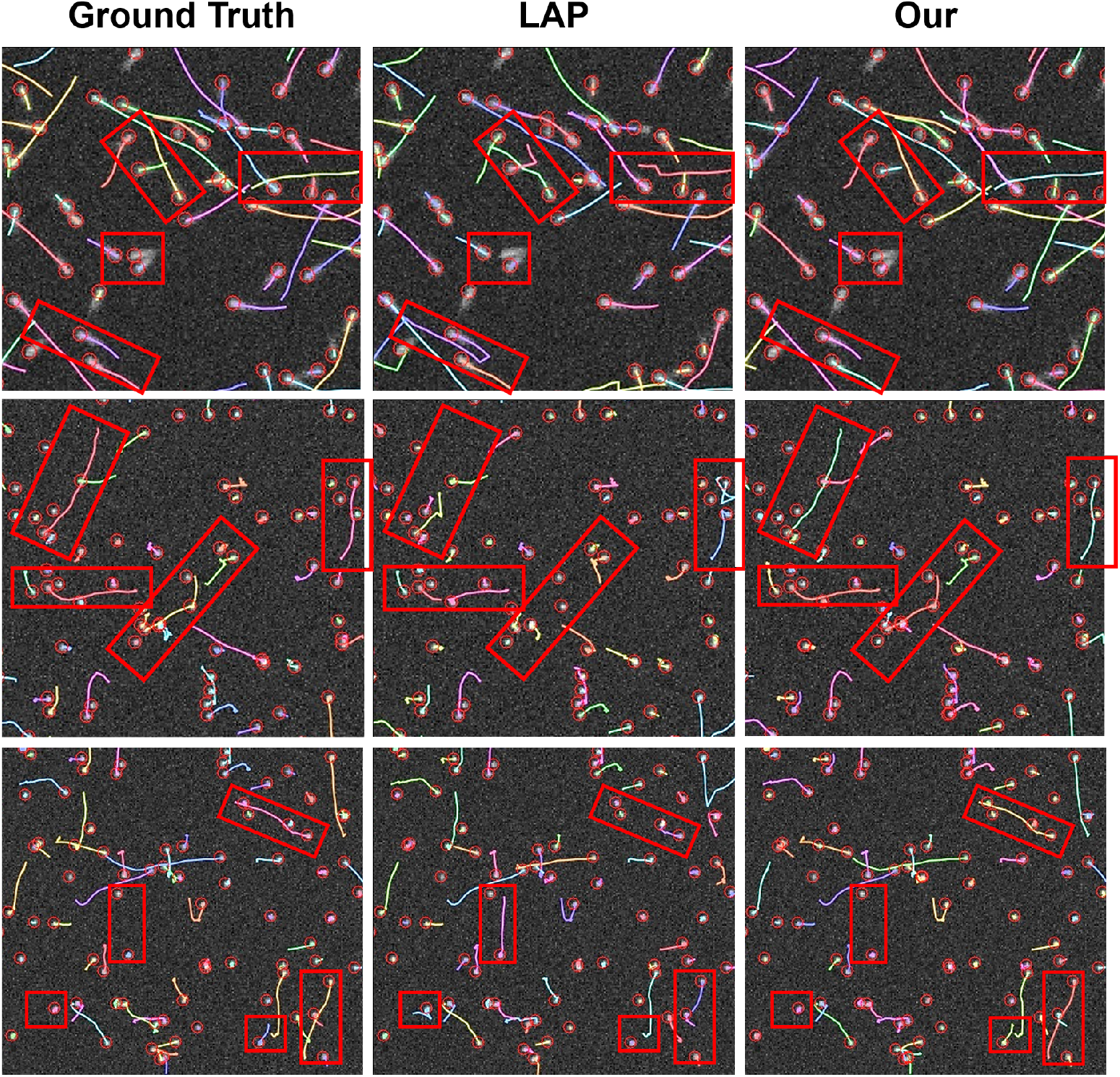
Particle Tracking performance comparison.

## Appendix F Visualization of the re-linking strategy effectiveness

**Fig. F6.**
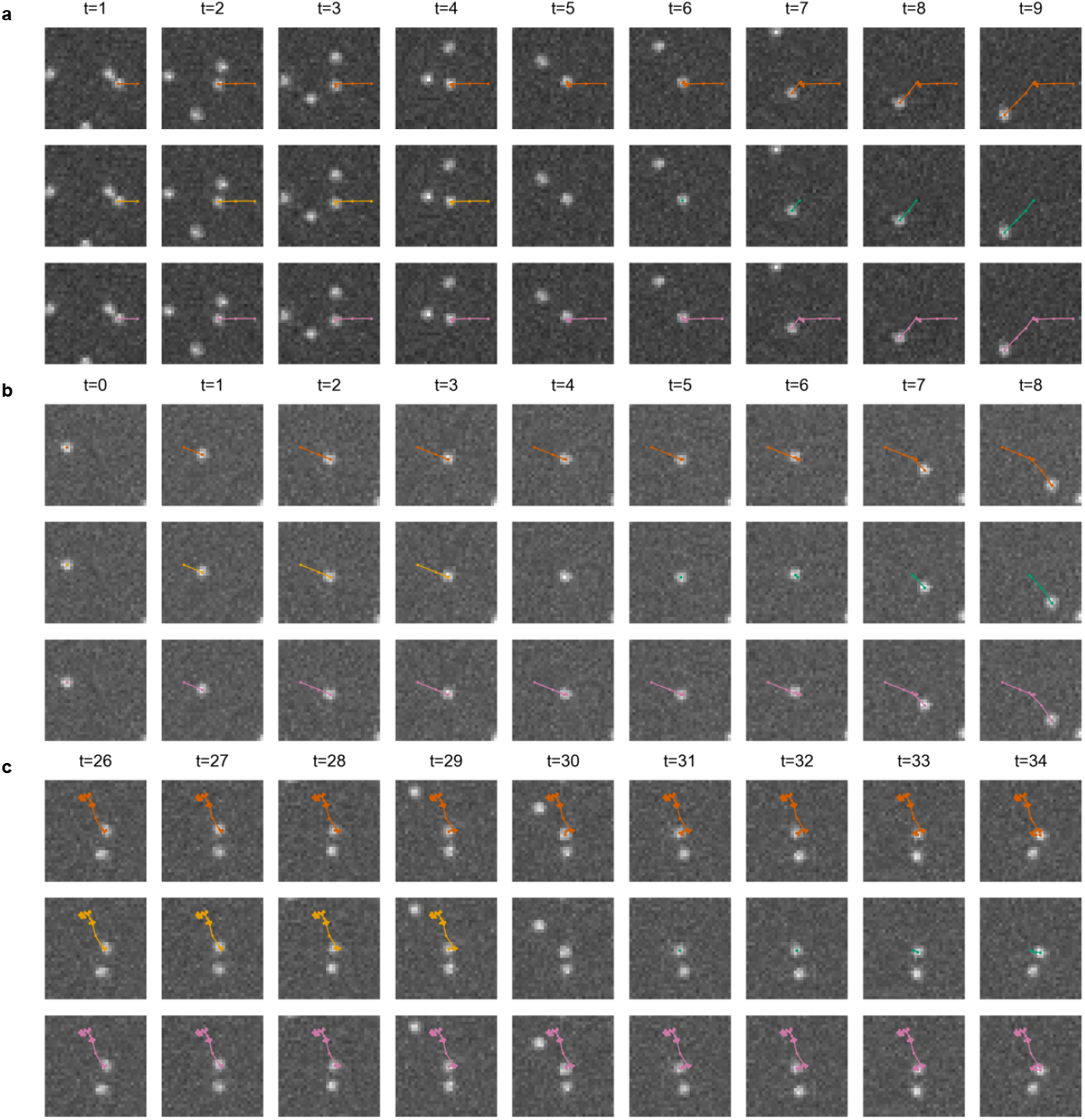
Visualization of the re-linking strategy effectiveness. **a-c**, Three examples under receptor scenario. The first row represents the ground truth trajectories, the second row represents the trajectories without the re-linking strategy, and the third row shows the trajectories with the re-linking strategy applied. Each example displays 9 consecutive frames. **a**, False negative occurs at frame t=5. **b**, False negative occurs at frame t=4. **c**, False negative occurs at frame t=30. The results indicate that without the re-linking strategy, false negatives cause a switch in track IDs, whereas the re-linking strategy maintains trajectory continuity, aligning more closely with the ground truth trajectories.

## Appendix G Management of tracks

### Algorithm 1

Management of tracks

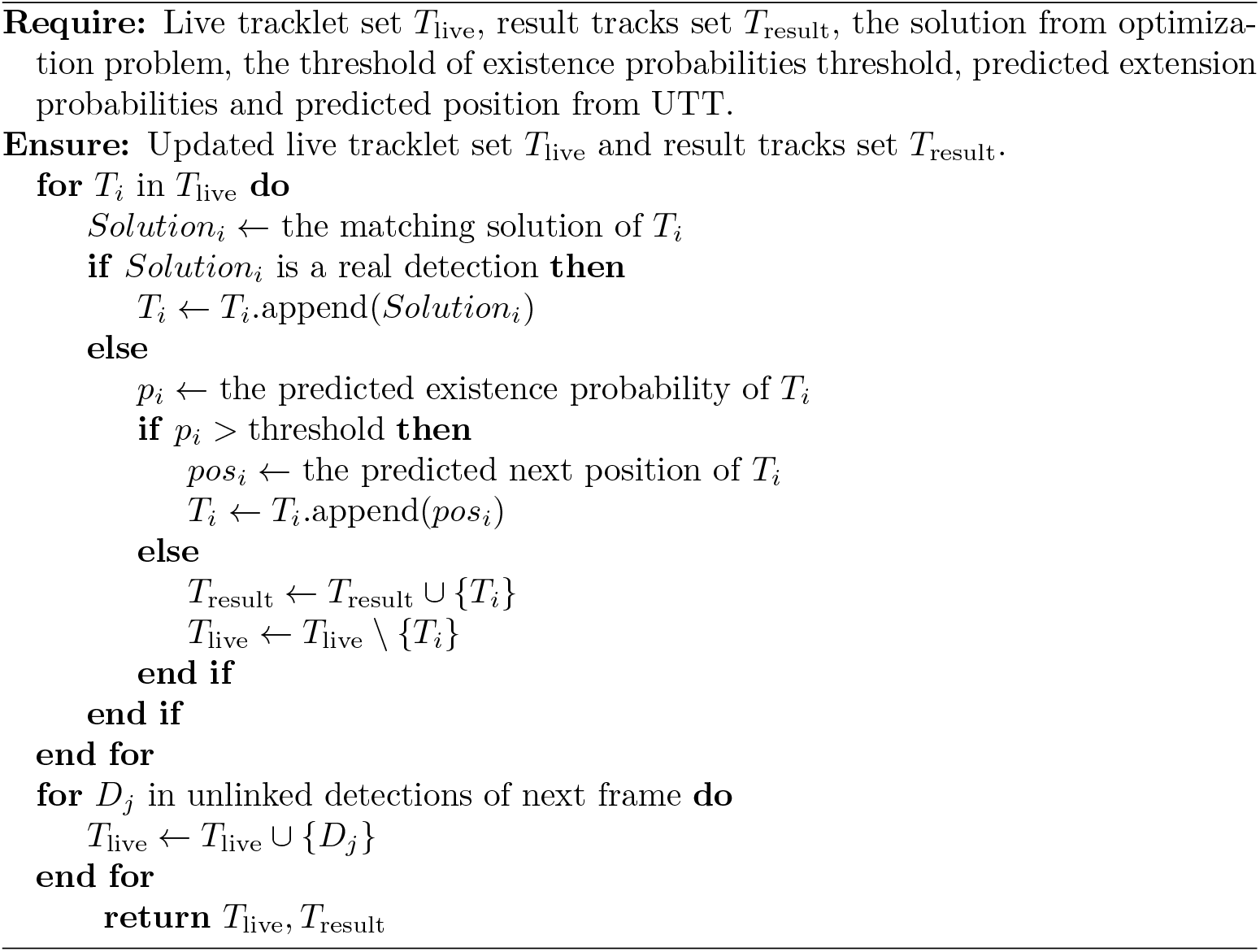

## Appendix H Lineage Reconstruction Algorithm

### Algorithm 2

Lineage Reconstruction Algorithm

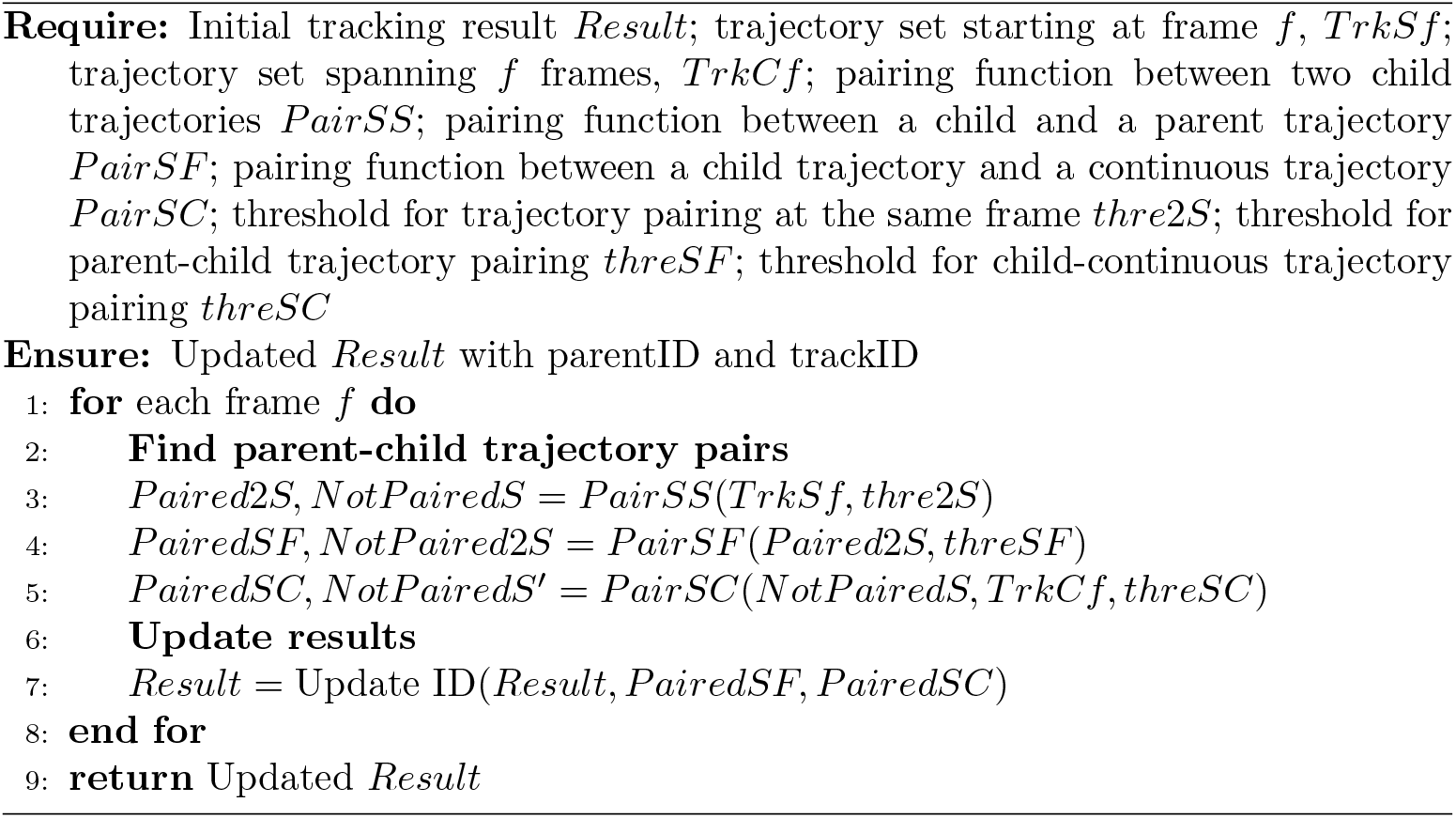

## Appendix I Software

**Fig. I7.**
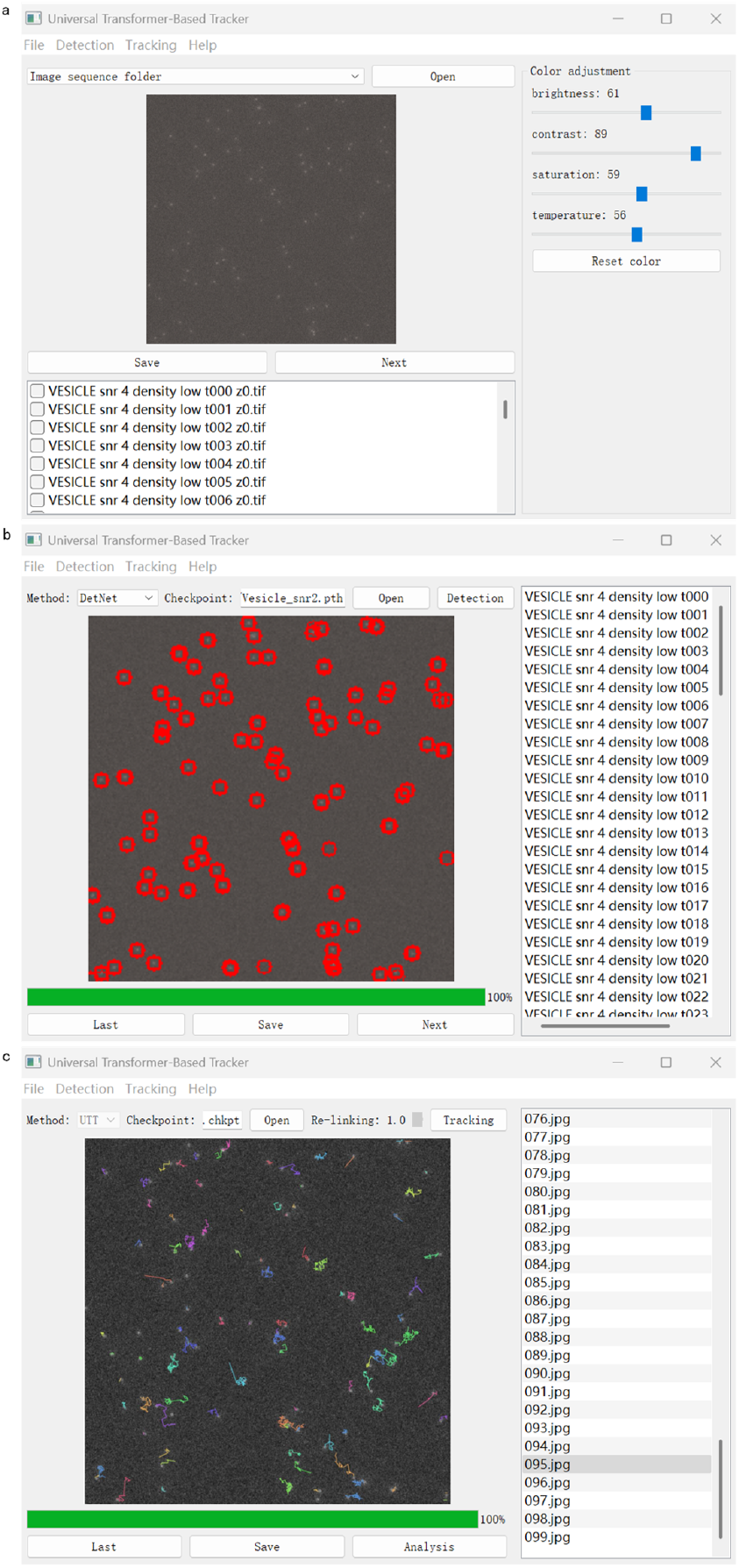
Graphical user interface of Universal Transformer-based Tracker. **a**, The file interface provides the following functions: opening image sequences folder and adjusting the image color. **b**, The particle detection interface provides the following functions: selecting detection model, labeling particles, and recording the particle center position. **c**, The tracking interface provides the following functions: loading pretrained detection model, configuring parameters, and generating particle trajectories.

